# The FAM114A proteins are adaptors for the recycling of Golgi enzymes

**DOI:** 10.1101/2024.03.27.587010

**Authors:** Lawrence G. Welch, Nadine Muschalik, Sean Munro

## Abstract

The Golgi apparatus contains many resident enzymes that must remain in place whilst their substrates flow through on their journey from the endoplasmic reticulum to elsewhere in the cell. COPI-coated vesicles bud from the rims of the Golgi stack to recycle Golgi residents to earlier cisternae. Different enzymes are present in different parts of the stack, and at least one COPI adaptor protein, GOLPH3, has been shown to recruit enzymes into vesicles in a specific part of the stack. We have used proximity biotinylation to identify further components of intra-Golgi transport vesicles and found FAM114A2, an uncharacterised cytosolic protein. Affinity chromatography with FAM114A2, and its paralogue FAM114A1 showed that they bind to numerous Golgi resident proteins, with membrane-proximal basic residues in the cytoplasmic tail being sufficient for the interaction. Deletion of both proteins from U2OS cells did not result in substantial defects in Golgi function. However, a *Drosophila* orthologue of these proteins (CG9590/FAM114A) is also localised to the Golgi and binds directly to COPI. Generation of *Drosophila* mutants lacking FAM114A revealed defects in glycosylation of glue proteins in the salivary gland. Thus, the FAM114A proteins are COPI vesicle resident proteins that bind to Golgi enzymes and so are candidate adaptors to contribute specificity to COPI vesicle recycling in the Golgi stack.

## INTRODUCTION

The Golgi apparatus is the major sorting hub in the secretory pathway. It receives newly made lipids and proteins from the endoplasmic reticulum (ER) and then sorts them to the cell surface or the compartments of the endocytic system. Following arrival from the ER, proteins and lipids move through the stack of Golgi cisternae before leaving in carriers forming at the trans side of the Golgi. The Golgi stack contains many resident enzymes that modify glycoproteins and glycolipids as they move through the stack (Moremen et al., 2012; Schjoldager et al., 2020). These enzymes, along with the transporters that deliver nucleoside-sugars and ions, must all maintain their residence within the stack whilst their substrates arrive and then depart. Furthermore, the enzymes are typically arranged within the stack in the order in which they act and so are localised to only a subset of cisternae. There has been much debate about the mechanism by which cargo proteins move past the enzymes that modify them, but the current widespread consensus is that the cisternae form on the cis side and then mature as they progress through the stack (Glick and Nakano, 2009; Pantazopoulou and Glick, 2019). As the cisternae progress, the resident enzymes are continuously recycled in vesicles and delivered to earlier cisternae in the stack and so maintain a constant distribution in a manner analogous to hopping down an upward moving escalator (Lujan and Campelo, 2021; Welch and Munro, 2019).

Vesicle budding from the Golgi stack is dependent on the COPI coat that is formed from the small GTPase Arf and coatomer, a heptameric complex distantly related to the clathrin adaptors proteins (Arakel and Schwappach, 2018; Beck et al., 2009; Taylor et al., 2022). COPI is responsible for the recycling of ER residents back to the ER from the cis Golgi, and it can bind directly to recycling signals in the cytoplasmic tails of ER membrane proteins. However, COPI is also present on later cisternae in the stack and in vitro budding assays have shown that it can concentrate Golgi resident proteins into vesicles (Adolf et al., 2019; Eckert et al., 2014). This raises two questions: how COPI can collect different cargo in different parts of the stack, and how are these vesicles then delivered to distinct destinations depending on where they originated from. In vertebrates, two of the coatomer subunits are present as pairs of paralogues, but in vitro vesicle budding assays did not detect significant differences between the contents of vesicles formed using coatomer containing different combinations of these paralogues (Adolf et al., 2019). Beyond the coat itself, another factor that could influence cargo recruitment is adaptors that bind specific subsets of cargo as is the case for clathrin-coated vesicles (Traub and Bonifacino, 2013). For COPI, the clearest example of such an adaptor is the cytosolic protein GOLPH3 which binds directly to the short basic tails found on many Golgi glycosylation enzymes (Ali et al., 2012; Schmitz et al., 2008; Tu et al., 2008; Welch and Munro, 2019). Removal of GOLPH3 results in loss of Golgi retention for a subset of Golgi residents, and addition of GOLPH3 to an in vitro COPI budding assay stimulates the incorporation of specific cargo into vesicles (Eckert et al., 2014; Rizzo et al., 2021; Welch et al., 2021).

Given that the Golgi contains a diverse population of resident proteins that varies between cell types it seems likely that there are additional adaptors that enable the COPI coat to select specific cargo in specific circumstances. To seek such adaptors, we have made use of one of the mechanisms by which the vesicles that bud from the Golgi are captured by their destination compartments. Capture of vesicles within the stack has been shown to depend, at least in part, on long coiled-coil proteins called golgins, with different golgins found in different parts of the stack (Gillingham and Munro, 2016; Muschalik and Munro, 2018; Witkos and Lowe, 2017). When these golgins are relocated to an ectopic location they are sufficient to cause ectopic vesicle capture, and different golgins capture different classes of vesicle (Park et al., 2022; Wong and Munro, 2014). Three golgins have been shown to capture Golgi-derived vesicles at an ectopic location: GMAP-210, golgin-84 and TMF, encoded in humans by the genes TRIP11, GOLGA5 and TMF1. GMAP-210 and golgin-84 are located on the cis/medial Golgi whilst TMF is later in the stack, and consistent with this, the vesicles captured by TMF contain proteins from the later part of Golgi (Bascom et al., 1999; Mori and Kato, 2002; Sato et al., 2015; Wong and Munro, 2014). The mechanism by which these golgins capture vesicles remains unclear, although it is known that motifs at the N-terminus are sufficient, and in the case of GMAP-210 it has been suggested that this region recognises the lipid composition of the vesicle (Magdeleine et al., 2016; Wong et al., 2017). Nonetheless, the ectopic capture of vesicles by golgins provides a means to isolate them from the rest of the stack and examine their contents. This has been successfully applied to the golgins at the trans-Golgi which capture carriers coming from endosomes and has allowed the identification of vesicle resident proteins and a linker protein that connects the golgins to the vesicles (Shin et al., 2017; Shin et al., 2020). In this paper we use the golgin GMAP-210 to identify the FAM114A proteins as being associated with intra-Golgi transport vesicles and demonstrate that they can bind Golgi residents and are required for normal Golgi function in vivo.

## RESULTS

### Identification of the FAM114A proteins as residents of intra-Golgi transport vesicles

To identify novel components of intra-Golgi vesicles, a MitoID proximity-dependent labelling assay was applied to the intra-Golgi golgin tether GMAP-210 as done previously for trans-Golgi golgins and Rab GTPases. In short, the basis of the screen was to ectopically relocate intra-Golgi vesicles to the mitochondria and biotinylate the proximal vesicle-resident proteins to allow purification by streptavidin pulldown and identification by mass spectrometry (Fig. 1A). To do this, the N-terminal vesicle-binding region of GMAP-210 was fused to the promiscuous biotin ligase BirA*, the coiled-coil protein GCC185 as a spacer that lacks tethering activity, an HA epitope tag and the mitochondrial targeting sequence of monoamine oxidase (MAO). Controls included a version where a conserved tryptophan residue is mutated as this has been shown to cause GMAP-210 to lose the ability to capture giantin- and GALNT2-containing vesicles (Wong et al., 2017). These golgin chimeras were stably expressed in HEK293 cells, and to capture vesicles, expression was induced, and cells treated with nocodazole to convert the Golgi into ministacks. Previous work has shown that this treatment increases the efficiency of capture of Golgi-derived vesicles as it places many more mitochondria in the proximity of a Golgi stack (Wong and Munro, 2014). Following a biotin pulse, labelled proteins were isolated with streptavidin and identified by mass-spectrometry.

**Figure 1.**
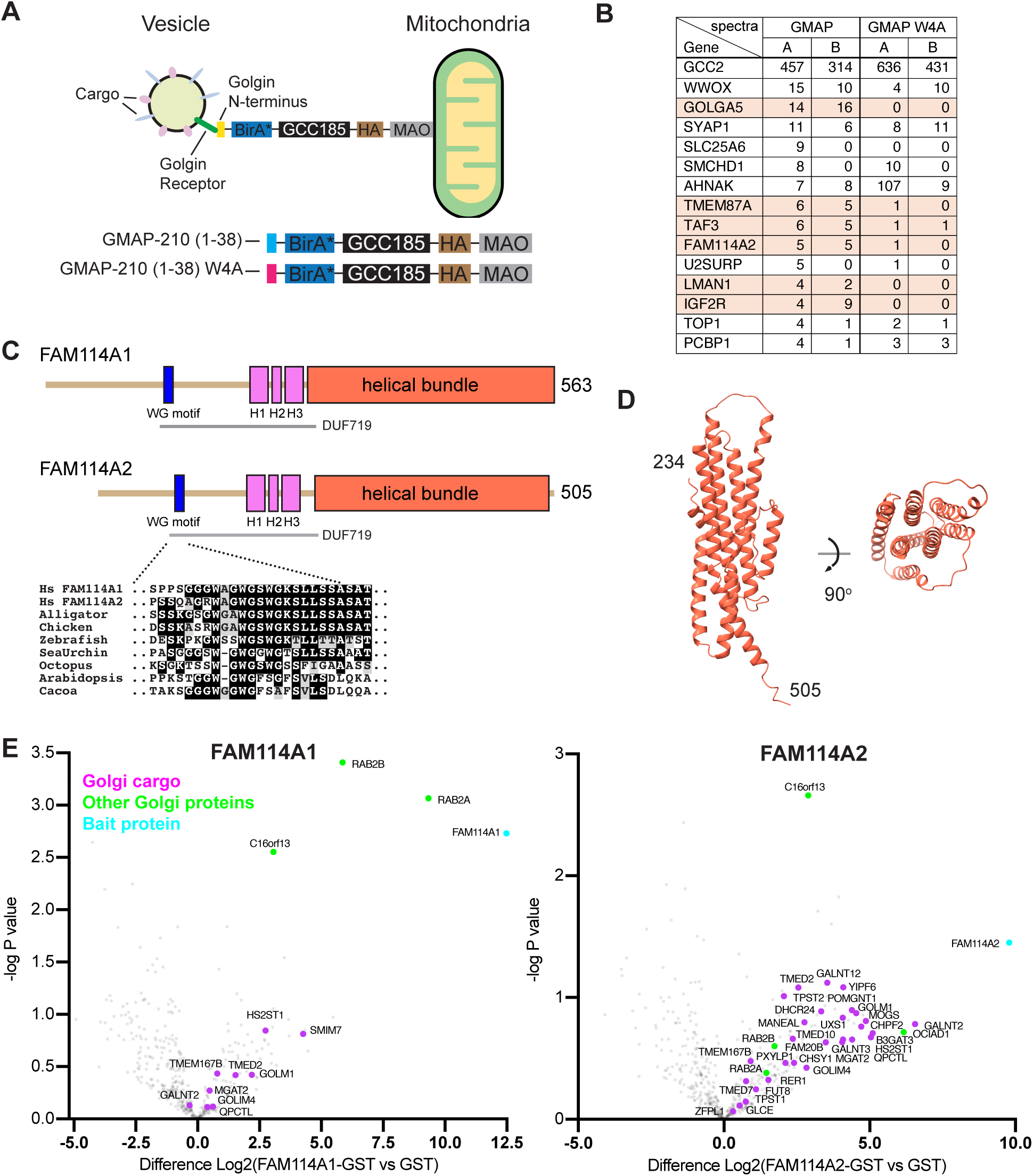
A MitoID screen identifies the FAM114A proteins as novel early-Golgi COPI vesicle- residents. **(A)** A schematic outlining the MitoID screen: the vesicle binding regions of intra-Golgi golgins were fused to BirA* and the large coiled-coil protein GCC185 and relocated to the mitochondria in Flp-In T-REx HEK293T cells. Ectopically rerouted intra-Golgi vesicles were subject to proximity-dependent biotinylation allowing for vesicle-resident proteins to be purified from cell lysate with streptavidin and identified by mass spectrometry. **(B)** The top 15 hits from the MitoID data that gave spectra with GMAP-210 (1-38) but no spectra with a negative control (Flp-In T-REx cells integrated with an empty pcDNA5/FRT/TO vector). Total spectral counts from two independent biological replicates are shown, ranked by GMAP-210 (B). Orange shading indicates hits that are found in both repeats and substantially reduced by the W4A mutation in the GMAP-210 vesicle capture motif. **(C)** The domains of FAM114A proteins based on AlphaFold2 predictions (Fig S1A). The N-terminal halves are predicted to be unstructured apart from three adjacent helical regions (H1-H3). There is also a highly conserved motif rich in tryptophan and glycine residues (WG), and a MUSCLE sequence alignment shows this region from diverse species. The region encompassing the WG motif and H1-H3 was designated a domain of unknown function by the Pfam database (DUF719). The C-terminal half of the proteins is predicted to form a bundle of seven helices. **(D)** AlphaFold2 prediction of the helical bundle formed by the C-terminal region of human FAM114A2. **(E)** Volcano plots showing the spectral intensities of proteins pulled down from HEK293T cell lysate using GST-tagged FAM114A proteins compared to GST alone. -log P values were generated from Welch’s t-tests. Indicated are Golgi-resident integral (magenta) and peripheral (green) membrane proteins (Swiss-Prot database) and bait proteins (cyan). Data represents three biological replicates.

Of the top 15 proteins enriched with the wild-type GMAP-210 bait, several known Golgi-resident proteins were identified including ERGIC-53 (LMAN1), golgin-84 (GOLGA5), TMEM87A and FAM114A2, all of which were substantially reduced by the W4A mutation in GMAP-210 vesicle capture motif (Fig. 1B). ERGIC-53 and golgin-84 are integral membrane proteins that are well characterised as cargo of Golgi-derived vesicles (Adolf et al., 2019). In contrast, FAM114A2 is a cytosolic protein of unknown function. The *Drosophila* orthologue, CG9590, was identified as a Rab2 effector in an affinity chromatography proteomic screen using S2 cell lysate, and this was subsequently supported by findings in a Rab MitoID screen in mammalian cells (Gillingham et al., 2014; Gillingham et al., 2019). The *Drosophila* protein and human FAM114A1 were both shown to colocalise with cis-Golgi markers, but their function is unknown. All are predicted by AlphaFold2 to have an unstructured N-terminal half and a C-terminal helical bundle (Fig. 1C,D; Fig. S1A). The unstructured N-terminus contains a highly conserved motif containing three tryptophan-glycine pairs. This motif is part of a region previously assigned as a Domain of Unknown Function (DUF719) by the Pfam database, but its role is unclear. The C-terminal helical bundle is predicted by AlphaFold to be the part that binds Rab2 (Fig. S1B).

### FAM114A proteins bind Rab2 and Golgi resident membrane proteins

To elucidate the function of the FAM114A proteins, GST-tagged forms of both were produced in bacteria and used as baits in pulldowns to isolate binding partners from HEK293T cell lysate. The interacting proteins were identified by mass spectrometry and compared to a control of GST alone using volcano plots. Both FAM114A proteins enriched Rab2A and Rab2B, and a selection of Golgi-resident integral membrane proteins (Fig. 1E). They also efficiently enriched METTL26 (C16orf13), a cytosolic protein of unknown function. However, this was not found as an interactor with the *Drosophila* orthologue (see below) and shows no predicted interaction with FAM114A1 or FAM114A2 using AlphaFold2 and so was not investigated further. FAM114A1 pulled down relatively few proteins whereas FAM114A2 specifically enriched a vast array of Golgi-resident membrane proteins, many of which are presumptive intra-Golgi COPI cargo proteins such as glycosyltransferases.

### FAM114A2 interacts directly with the tails of Golgi enzymes through their membrane-proximal poly-basic residues

FAM114A proteins are predicted to be peripheral membrane proteins and so it is likely that they are binding the cytoplasmic tails of Golgi resident proteins. This is reminiscent of the COPI adaptors GOLPH3 and GOLPH3L, which have been shown to interact with the cytoplasmic tails of Golgi residents through membrane-proximal polybasic stretches (Ali et al., 2012; Rizzo et al., 2021; Tu et al., 2008; Welch et al., 2021). The predicted structure of the FAM114A helical bundle also has large electronegative surfaces like the surface observed on the crystal structure of GOLPH3 (Fig. S1C). To examine the interaction with Golgi enzymes, we applied to FAM114A proteins a binding analysis like that applied to GOLPH3 (Welch et al., 2021). In this assay, the signal anchor region of a plasma membrane protein, sucrase-isomaltase (SI), is expressed as a fusion to GFP, and the cytoplasmic tail replaced with those of various type II Golgi-resident proteins (Fig. 2A). The fusion proteins are expressed in mammalian cells and after solubilisation in detergent their binding to FAM114A-coated beads can be assayed. We initially tested direct binding of FAM114A proteins to the tail of GALNT2, a Golgi-resident O-linked mucin type glycosyltransferase which was pulled down by FAM114A2. The GALNT2-SI-GFP-FLAG chimera was recombinantly expressed and purified from HEK293T cell lysate using FLAG affinity chromatography. The purified chimera was then assayed for binding to beads coated with GST-tagged FAM114A proteins. The GALNT2 tail chimera exhibited strong, direct and specific binding to GST-tagged FAM114A2 but not to GST-tagged FAM114A1 or GST alone (Fig. 2B). Next, we generated a series of chimeras with tails from a range of different Golgi enzymes and tested the ability of GST-tagged FAM114A2 to pull them out from HEK293T cell lysates, as we had done for GOLPH3 (Welch et al., 2021). As with GOLPH3, FAM114A2 bound to diverse tails with a preference for tails with membrane-proximal polybasic clusters. In contrast, tails with a paucity of positive residues or those containing negative residues, such as the plasma membrane protein SI, bound poorly to FAM114A2 (Fig. 2C). Membrane proximal insertion of 3 arginines or lysines, but not histidines (which are not protonated at a cytosolic pH of 7.4), into the tail of SI was sufficient to bestow FAM114A2 binding. In summary, FAM114A2 is comparable to the COPI adaptors GOLPH3 and GOLPH3L as it binds directly to the tails of cargo with membrane-proximal polybasic stretches.

**Figure 2.**
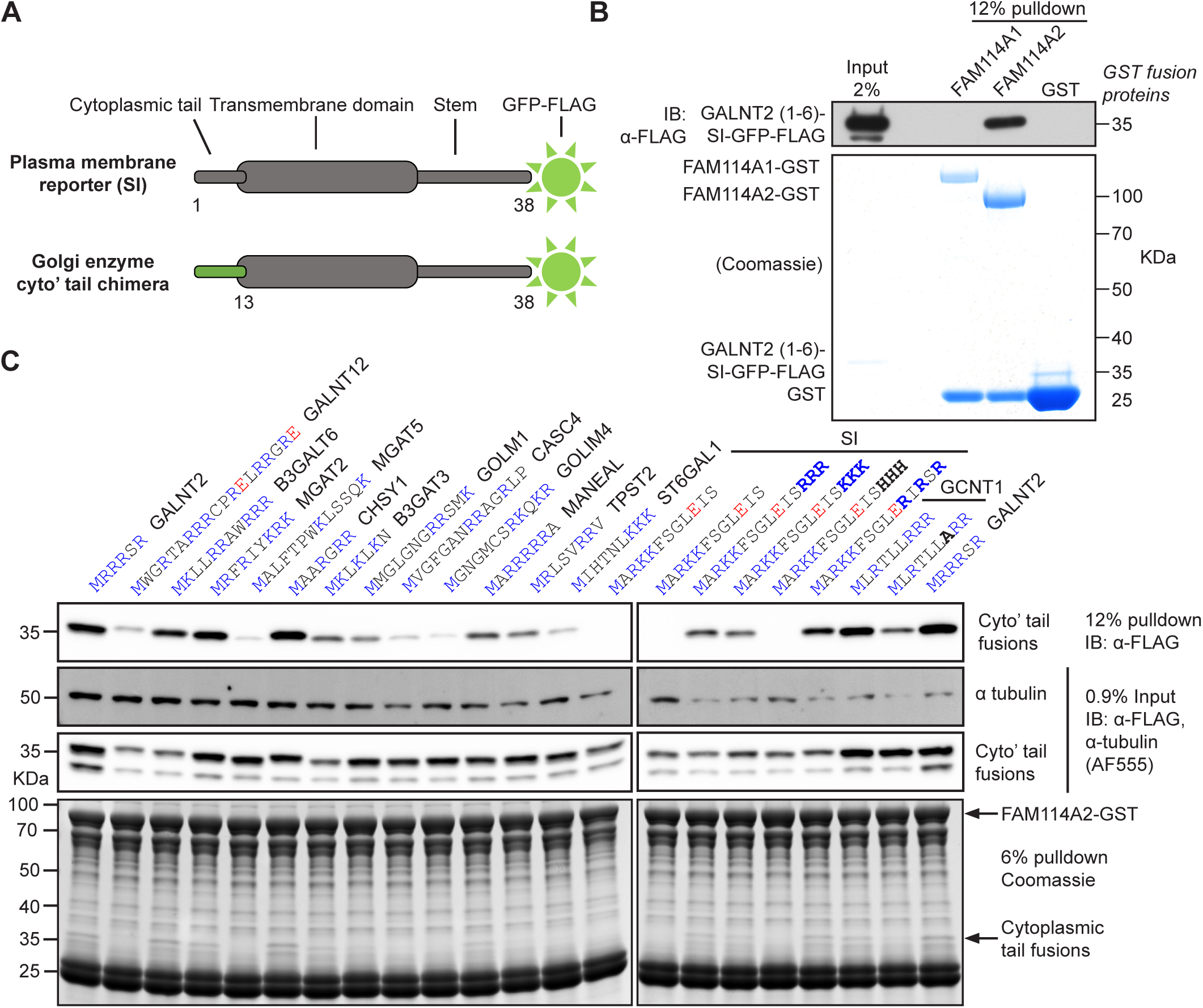
FAM114A2 binds directly to the tails of cargo with membrane-proximal poly-basic stretches. **(A)** A cartoon schematic of the GFP-tagged cytoplasmic tail chimeras used in binding studies. Sucrase-isomaltase (SI). **(B)** A GST pulldown showing direct binding of FAM114A2 to the tail of GALNT2. GST-tagged FAM114A proteins and GST were expressed in Sf9 cells using baculoviral transduction while the GALNT2 chimera was expressed in HEK293T cells. The GST-tagged baits and GALNT2 chimera were independently purified from lysates using GST and FLAG affinity chromatography respectively. The GALNT2 chimera was eluted using excess FLAG peptide prior to mixing with GST bait-loaded beads. **(C)** Binding studies to test the ability of FAM114A2-GST to pulldown different cytoplasmic tail chimeras from HEK293T cell lysate. Tail sequences and their corresponding gene names (above) are coloured in blue (positive) or red (negative) according to their predicted charge at a cytosolic pH of 7.4. Data is representative of two independent experiments.

### Effects of deletion of FAM114A genes from cultured cell lines

Deletion of GOLPH3 and GOLPH3L from U2OS cells leads to reduced levels of a subset of Golgi enzymes and downstream defects in glycosylation (Welch et al., 2021). Thus, CRISPR-Cas9 gene-editing was used to delete the FAM114A genes in wild-type U2OS cells and also in *ΔΔGOLPH3, GOLPH3L* U2OS cells in case of functional redundancy between the GOLPH3 and FAM114A families (Fig. 3A). Multiplexed quantitative mass spectrometry was used to compare relative protein abundances between wild-type and *ΔΔFAM114A1, FAM114A2* U2OS cell lines. In contrast to what was seen with *ΔΔGOLPH3, GOLPH3L* cells, there was no clear difference in the levels of Golgi-resident proteins in wild-type U2OS cell vs *ΔΔFAM114A1, FAM114A2* cells (Fig. S2A). We had previously shown that the deletion of GOLPH3 genes perturbed the Golgi-retention of a GALNT2 cytoplasmic chimera in an *in vivo* Golgi retention assay (Welch et al., 2021). We found that deletion of both FAM114A genes in a wild-type U2OS background had a far smaller effect on Golgi retention of the GALNT2 reporter when analysed by flow cytometry or immunofluorescence (Fig. 3B; Fig. S2B). Furthermore, there was no detectable additive effect when the FAM114A genes were deleted in a GOLPH3 double knockout background. As reported previously, a panel of cell surface lectins revealed strong defects in glycosylation in the *ΔΔGOLPH3, GOLPH3L* U2OS cells, but in contrast knockout of the FAM114A genes did not result in consistent changes in lectin labelling apart from a small increase in binding by WFA which recognises mucin-type O-linked glycans (Fig. S2C).

**Figure 3.**
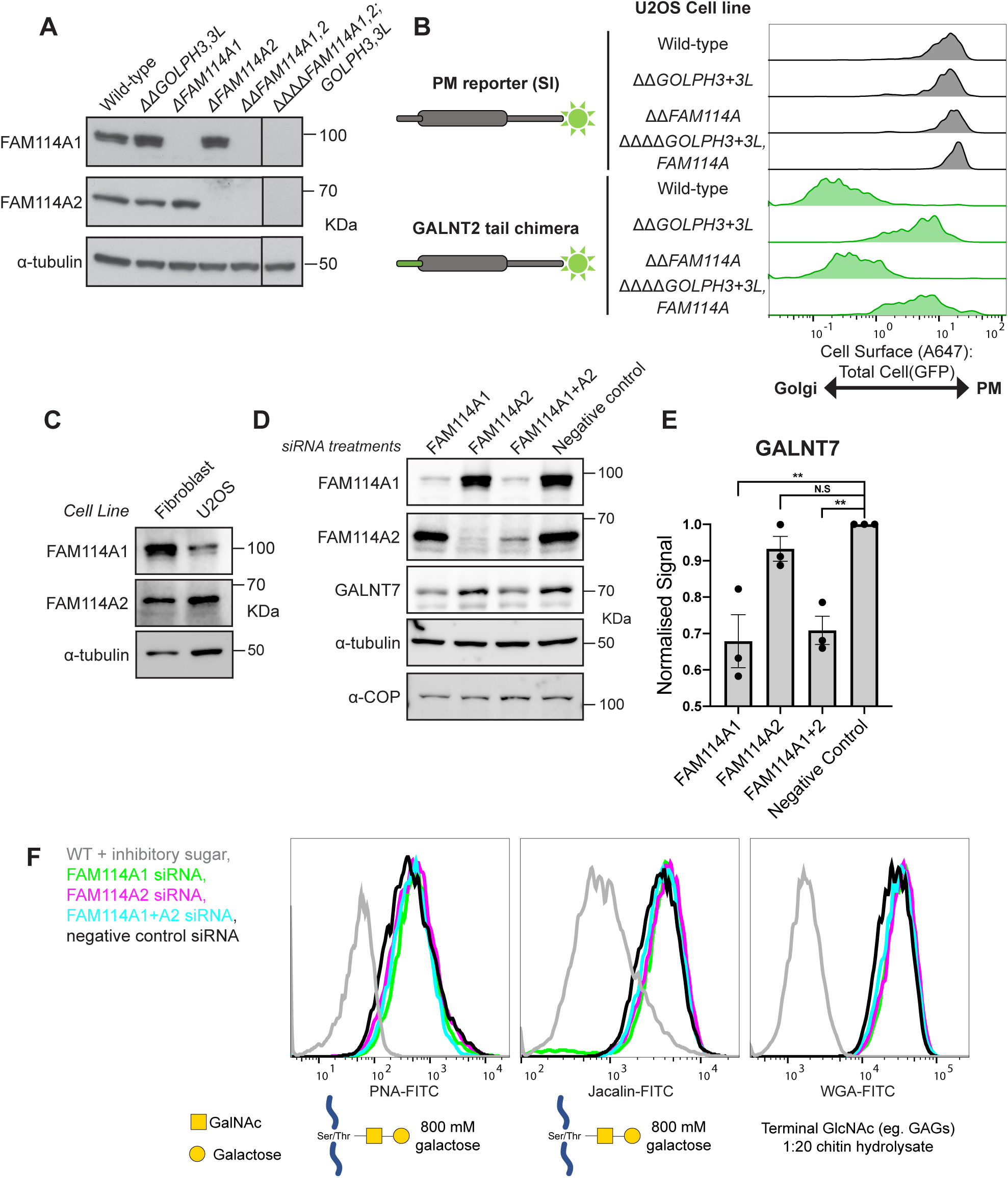
Disruption of FAM114A genes in U2OS cells and primary skin fibroblasts. (A) Immunoblots of FAM114A and GOLPH3 family combinatorial U2OS CRISPR-Cas9 knockout cell lines. For validation of disruption of the *GOLPH3* and *GOLPH3L* loci, see Welch et al., 2021. (B) Histograms from an *in vivo* Golgi retention assay comparing the retention of different reporters in various U2OS knockout cell lines. Histograms represent a ratio of the A647 cell surface signal and the GFP total cell signal of the reporters and therefore serve as a quantitative readout for Golgi retention. Sucrase-isomaltase (SI). Histograms represent approximately 500 events and are representative of 3 biological replicates. (C) Immunoblots comparing the levels of FAM114A proteins in human skin fibroblasts to U2OS cells. (D) Immunoblots comparing the levels of O-linked mucin-type glycosyltransferase GALNT7 after knockdown of FAM114A family members in fibroblasts. Negative control cells were treated with a non-targeting siRNA control. (E) Quantification of the difference in GALNT7 levels across different treatments as observed in (D). The integrated densities of the primary band for GALNT7 were normalised to that of the α-tubulin loading control which was then normalised to the negative control sample. Shown are mean normalised scores ± SEM from 3 independent experiments. Normalised scores were tested using a repeated measures one-way ANOVA with a Dunnett’s multiple comparisons test. **, P < 0.01. (F) Flow cytometry analysis of cell surface lectin-stained cells as treated in (D). A range of Lectins with specificities for different glycans (below) were used and were also applied in the presence of a saturating concentration of an inhibitory sugar to confirm the specificity of the stains. Histograms represent approximately 10,000 events.

It is possible that the FAM114A proteins are of more importance in specific cell types, and we also examined primary fibroblasts which express a different ratio of the FAM114A proteins than do U2OS cells (Fig. 3C). However, siRNA knockdown of the FAM114A proteins did not result in detectable changes in lectin staining, although we did detect a small but reproducible reduction in the levels of GALNT7, a Golgi enzyme whose level is also particularly sensitive to loss of GOLPH3 (Welch et al., 2021) (Fig 3D-F).

### The *Drosophila* FAM114A protein, CG9590, interacts with the COPI coat and Golgi resident proteins

For many proteins involved in Golgi function, their removal only causes detectable phenotypes in particular tissues, perhaps reflecting plasticity and robustness in intracellular trafficking pathways (Bem et al., 2011; Lowe, 2019; Marin-Valencia et al., 2017; Schmidt et al., 2007). Therefore, we used the *Drosophila* system to examine FAM114A activity in a multicellular organism. *Drosophila* have a single orthologue of FAM114A1/2, CG9590, which has previously been shown to bind to Rab2 but is otherwise uncharacterised (Gillingham et al., 2014). Expression atlas data shows that it is expressed in most tissues, with an elevation in cells with high secretory activity such as the salivary and accessory glands (Krause et al., 2022). The genes that match this tissue profile most closely are other proteins involved in Golgi function such as COPI subunits and SNAREs.

CG9590 is predicted to have a structure very similar to that of its mammalian orthologues, with an N-terminal unstructured domain including a region with WG motifs, and a C-terminal helical bundle with an electronegative surface (Fig. 4A). However, unlike the mammalian relatives, CG9590 also has near its N-terminus a motif containing two tryptophans embedded in acidic residues (^24^WDDW), and this feature is conserved amongst insects. A similar Wx_n(1-6)_[W/F] motif has been shown to bind to the μ-homology domain of the δ-COP subunit of coatomer and typically contains two tryptophan or phenylalanine residues separated by 2-3 residues and positioned within a highly acidic stretch (Suckling et al., 2015). As with the mammalian FAM114A proteins, we initially used affinity chromatography to identify potential binding partners of CG9590. CG9590 and for comparison the *Drosophila* orthologue of GOLPH3, Sauron, were expressed in bacteria as GST fusions and used to enrich interacting partners from S2 cell lysates. When compared to GST alone, GST-tagged CG9590 specifically enriched a plethora of membrane proteins that are resident in the Golgi (including several O-linked mucin-type glycosylation enzymes) or are likely to cycle between ER and Golgi, as well as various other trafficking components (Fig. 4B). As expected, Sauron also enriched Golgi residents, albeit with differing efficiencies compared to CG9590 (Fig. 4C). Neither protein showed an enrichment of the *Drosophila* ortholog of METTL26/C16orf13 (CG18661), the protein found enriched with the mammalian proteins (Figs. 4C and 4D). In addition to the lack of METTL26, another striking difference with the results with the human proteins was that CG9590 showed a strong enrichment of the subunits of the COPI coat. In order to determine if CG9590 was interacting with the COPI coat via the ^24^WDDW sequence that resembles a Wx_n(1-6)_[W/F] motif, the two tryptophans were mutated to alanine and the protein interactome of the mutant was compared to that of the wild-type protein. Relative to wild-type CG9590, the enrichment of the COPI coat subunits, Vap33 and subunits of the OST complex was markedly reduced in the mutant CG9590 sample (Fig. 4D). In contrast, there was little or no difference in binding to intra-Golgi proteins suggesting that the ability of CG9590 to bind Golgi residents is independent of COPI binding, whereas some ER residents are possibly binding directly to the COPI coat itself. Given the structural and functional similarities between the mammalian FAM114A proteins and *Drosophila* CG9590 we will refer to the latter as FAM114A.

**Figure 4.**
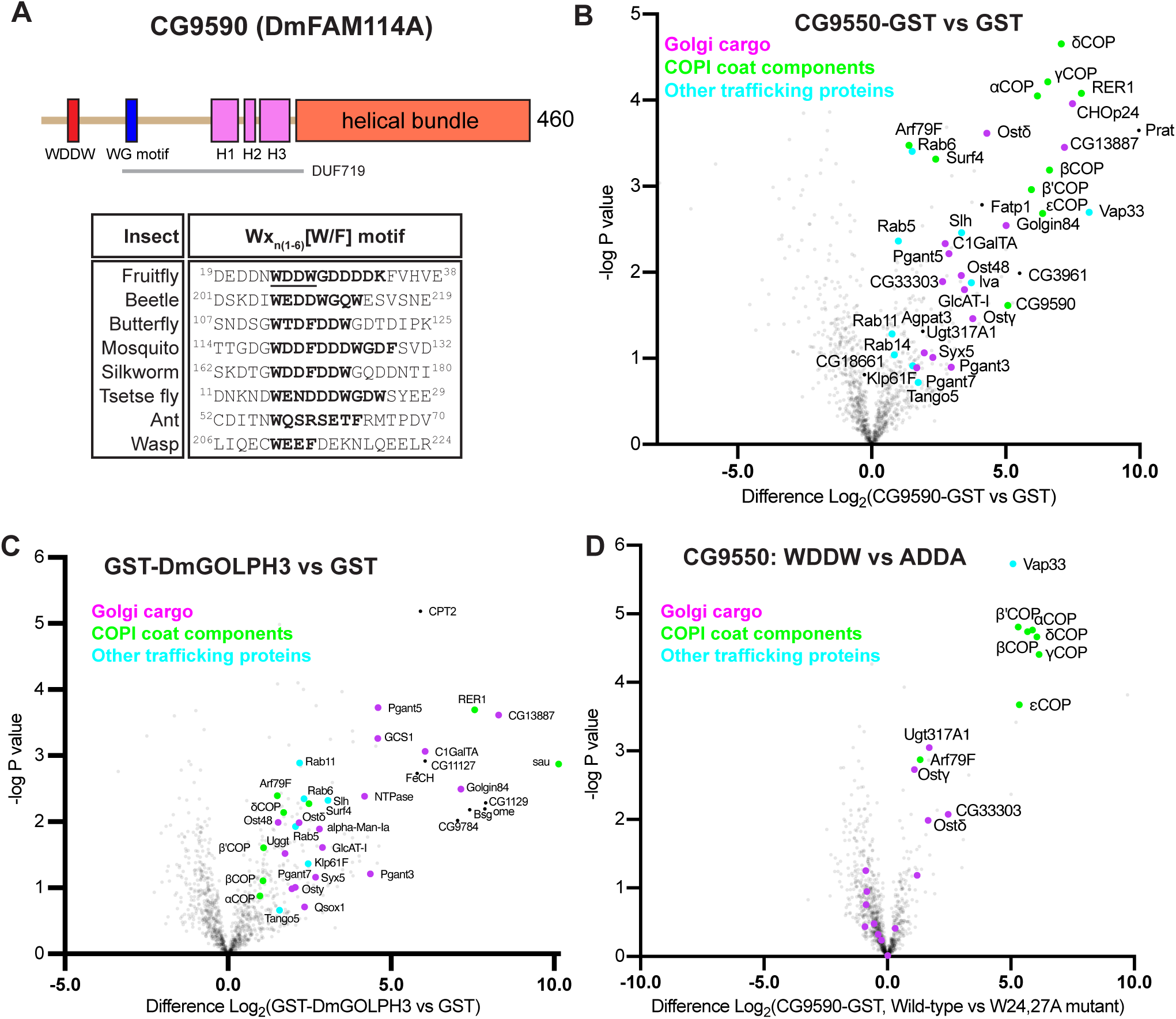
The *Drosophila* FAM114A ortholog, CG9590, binds to Golgi resident membrane proteins and the COPI coat. **(A)** A schematic of the different domains of CG9590, coloured and labelled as for the human proteins (Fig. 1C). In addition, insect orthologs share a conserved Wx_n(1-6)_[W/F] motif (WDDW) at their N-terminus which is predicted to bind the δ-COP subunit of the COPI coat. **(B)** Volcano plot showing the spectral intensities of proteins pulled down from S2 cell lysate using GST-tagged FAM114A compared to GST alone. -log P values were generated from Welch’s t-tests. Indicated are Golgi-resident integral membrane proteins (magenta), COPI coat components (green) and other known trafficking proteins (cyan). Data represents three biological replicates. **(C)** as (B) except using GST-tagged Sau (*Drosophila* GOLPH3). **(D)** as (B) except comparing GST-tagged FAM114A to the same fusion but with mutations in the two tryptophans in the putative coatomer binding region (W24A, W27A). These mutations primarily affect binding of coatomer, along with a few residents of the ER which may be binding to coatomer.

### Characterisation of a *Drosophila* mutant lacking FAM114A

To investigate the role of *Drosophila* FAM114A we used CRISPR-Cas9 to delete the entire gene from the genome (Fig. 5A). Flies lacking both alleles were viable and fertile, and an antibody raised against the *Drosophila* protein revealed that the protein was absent as expected. The antiserum was not suitable for immunofluorescence and so we generated *Drosophila* lines expressing a GFP-tagged form of FAM114A under UAS control (Fig. 5B). The tissue reported to have the highest level of expression of FAM114A is the larval salivary gland that secretes large amounts of glue proteins and has an abundance of secretory organelles and proteins (Loganathan et al., 2021). FAM114A-GFP was expressed in the salivary gland using a fkh-Gal4 driver and was found to accumulate on the Golgi apparatus (Fig 5C). Comparison to other Golgi markers showed that the protein is localised on the cis side of the Golgi stack being distributed between ER exit sites and the earliest Golgi markers such as the golgins GMAP and GM130. This is consistent with a role in recycling of Golgi residents or escaped ER residents from the earliest compartments of the stack.

**Figure 5.**
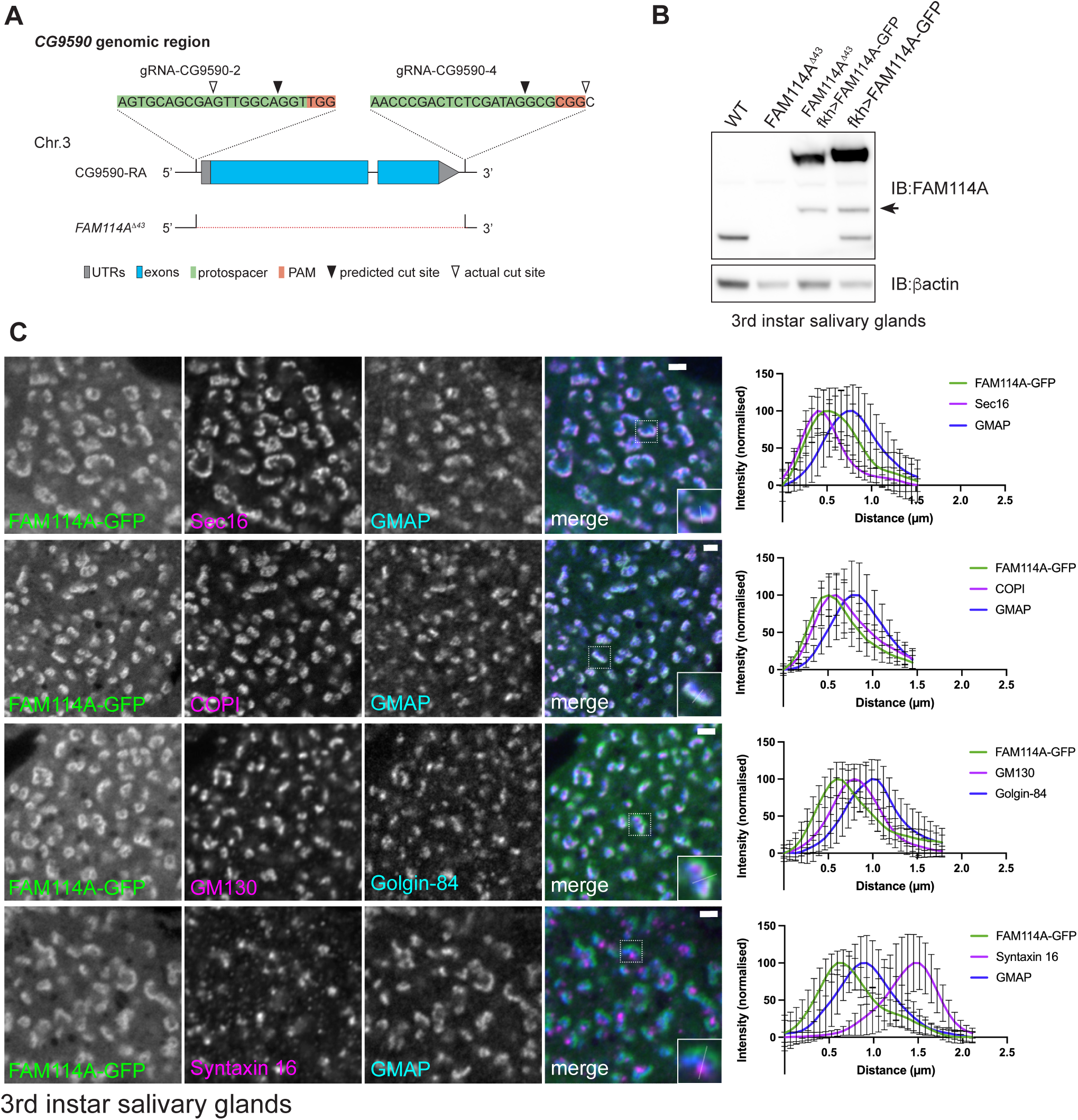
Drosophila FAM114A (CG9590) is localised to the Golgi. **(A)** Schematic of the *CG9590* genomic locus. A pair of gRNAs was used to delete the entire coding region including both UTRs. FAM114A^Δ43^ is a null allele that carries a deletion of 1812bp, deleting the entire coding region. **(B)** Immunoblot confirming the loss of FAM114A from 3^rd^ instar larval salivary glands in *FAM114A^Δ43^* mutants. UAS-FAM114A-GFP expressing flies were generated for localisation studies and rescue experiments. Expression of UAS-FAM114A-GFP using *forkhead(fkh)-Gal4* leads to elevated proteins levels when compared to wild type. FAM114A-GFP seems to be cleaved at low levels (arrow). **(C)** Confocal micrographs of 3^rd^ instar larval salivary glands cells expressing UAS-FAM114A-GFP using *fkh-Gal4* (green) labelled with various markers for ER exit sites and Golgi (magenta and blue). Line profiles were used for localisation analysis with graphs showing the mean of 10 line profiles across a Golgi stack. FAM114A-GFP seems to localise in between ER exit sites and the cis Golgi, and partially overlaps with coatomer (β’COP). Scale bars 2µm.

The glue proteins produced in the salivary gland include secretory mucins that are heavily modified with O-linked glycans and have thus proven useful for detecting defects in Golgi-dependent glycosylation (Biyasheva et al., 2001; Reynolds et al., 2019; Tran and ten Hagen, 2013). The major glue protein Sgs3 showed increased gel mobility in the *FAM114A* mutant and a deficiency removing the FAM114A gene, indicating reduced glycosylation, and this could be rescued by expression of FAM114A-GFP (Fig. 6A). The O-linked glycans that are attached to the glue proteins are initiated by the addition of GalNAc in the Golgi which is typically extended with galactose to form Galβ1,3GalNAc, and then extended further with glucuronic acid (Ji et al., 2018; Tian and ten Hagen, 2007). The lectin *Vicia villosa* agglutinin (VVA) recognises O-linked GalNAc and so labels the Golgi, and loss of this staining has been previously observed in mutants of the golgin coiled-coil proteins where Sgs3 mobility is also increased (Park et al., 2022). Loss of FAM114A results in smaller granules as shown by PNA lectin staining and loss of clear Golgi labelling by VVA (Fig. 6B). Taking these results together we conclude that FAM114A is required for normal Golgi glycosylation in *Drosophila*.

**Figure 6.**
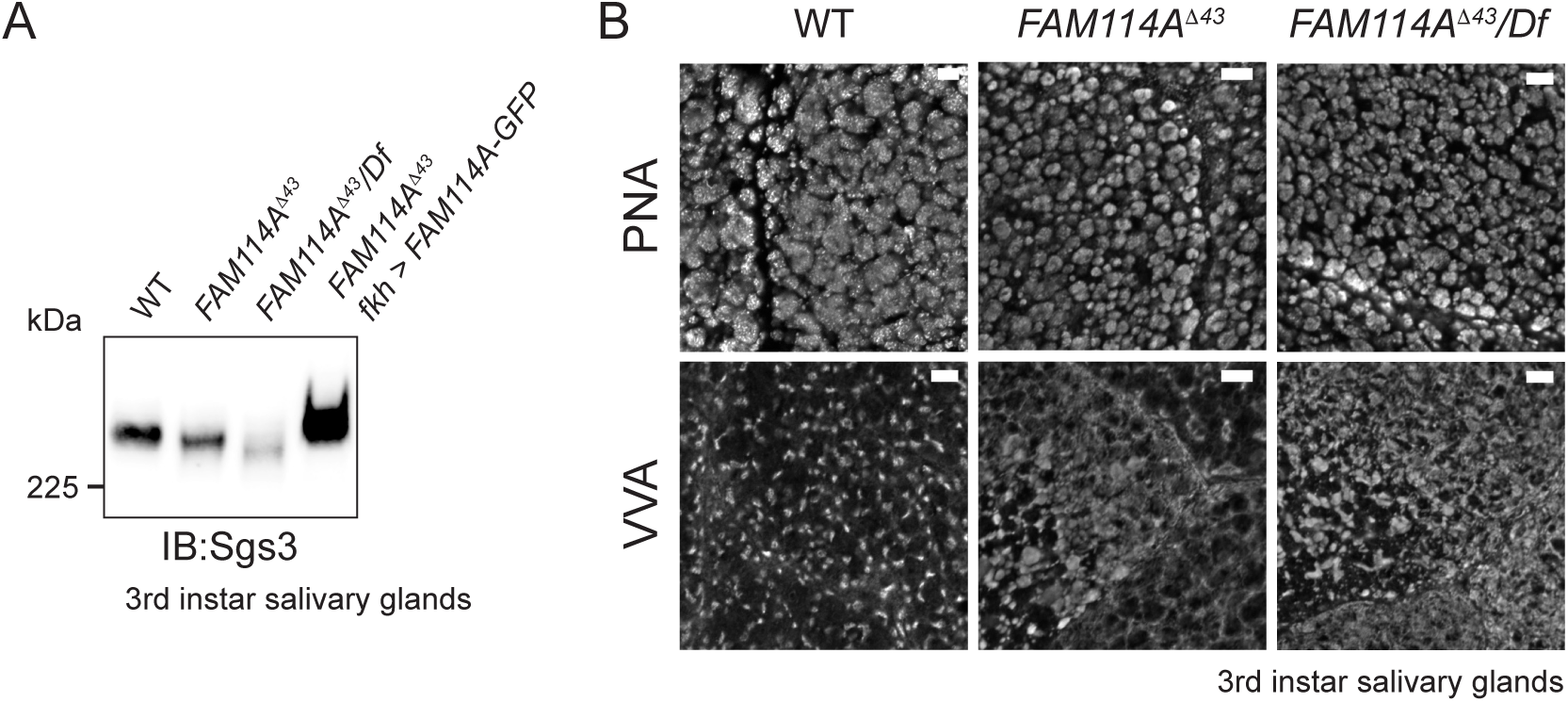
FAM114A mutants exhibit glycosylation defects. **(A)** Immunoblot of total protein extracts from 3^rd^ instar larval salivary glands from wild type, *FAM114A^Δ43^* mutants (homozygous and over a deficiency uncovering *CG9590*), and flies expressing the UAS-FAM114A-GFP rescue construct in the mutant background. The blot was probed for the heavily glycosylated mucin Sgs3. *FAM114A^Δ43^* mutants exhibit a mobility shift in Sgs3 which is rescued upon expression of FAM114A-GFP **(B)** Staining with lectins Peanut agglutinin (PNA) and *Vicia villosa* agglutinin (VVA) of 3^rd^ instar larval salivary gland cells of wild type, *FAM114A^Δ43^*mutants (homozygous and over a deficiency uncovering *CG9590*). PNA stains the secretory granules, which are smaller in *FAM114A* mutants. Lectin specificity was confirmed by use of the competing sugar *D*-galactose (Gal) at 0.3M. VVA stains the Golgi in wild type with specificity confirmed using the competing sugar N-acetyl-galactosamine (GalNAc) at 0.3 M. The VVA signal is nearly completely lost from the Golgi in FAM114A mutants. Scale bars 5µm.

## DISCUSSION

In this study we report that the FAM114A proteins are associated with the Golgi and intra-Golgi transport vesicles and that they can bind directly to the tails of Golgi resident enzymes via membrane-proximal basic residues. Removal of the single orthologue from *Drosophila* results in defects in glycosylation. From this we conclude that the FAM114A proteins act as adaptors to help recruit Golgi enzymes into COPI-coated vesicles that recycle membrane proteins within the Golgi stack and hence maintain the levels and organisation of glycosylation enzymes and other Golgi residents.

Such a role for the FAM114A proteins would be analogous to that of GOLPH3 which also binds to the tails of Golgi resident proteins and has been shown to promote their inclusion into COPI vesicles and to maintain their Golgi localisation (Rizzo et al., 2021; Tu et al., 2008; Welch et al., 2021). Unlike the FAM114A proteins, GOLPH3 is localised toward the trans side of the Golgi and thus is likely to function in a distinct set of COPI-coated vesicles. The FAM114A proteins appear to be more minor players in Golgi recycling than GOLPH3 given that more proteins are affected in cells lacking the latter, and that the GOLPH3 orthologue in *Drosophila* is essential for viability whereas FAM114A is not.

Individual knockouts for the two FAM114A paralogues in mice show no detectable effect, although a double mutant has not been reported (Khan et al., 2021; Muñoz-Fuentes et al., 2018). In contrast, GOLPH3 knockout in mice causes a range of severe phenotypes even though its less highly expressed paralogue GOLPH3L is still present (Muñoz-Fuentes et al., 2018). This seemingly more minor role for the FAM114A proteins is also suggested by it being absent from some invertebrates such as *C. elegans* and from the fungal phylum, whereas GOLPH3 is conserved in both. However, it is interesting to note that there appears to be a distant FAM114A orthologue in plants given that one enigma about GOLPH3 is that it is absent from plants despite them typically having well organised Golgi stacks populated by many different enzymes. In Arabidopsis this putative orthologue is encoded by the gene AT2G15860, with orthologues present in all plants examined, including green algae. For reasons that are obscure, it is annotated in UniProt as having a BAT2 domain, a term applied to the unstructured N-terminus of PRRC2 proteins, but such a domain is not detected by either the Pfam or InterPro domain databases. AlphaFold2 predicts a structure like that of the FAM114A proteins with an unstructured N-terminal region containing a tryptophan-rich motif and a C-terminal helical bundle. No functional characterisation has been reported in any plant species, but it seems a plausible candidate to have a role in the organisation of the Golgi in plants.

The FAM114A proteins are known to bind Rab2 in both humans and *Drosophila*. Our affinity chromatography with FAM114A1 and FAM114A2 found that the former binds most efficiently to Rab2 but this may reflect this in vitro assay system as an interaction between Rab2A and FAM114A2 was readily detected in our previous in vivo proximity biotinylation screen. AlphaFold2 gives a high confidence prediction for a complex between Rab2 and the FAM114A proteins, indicating that the interaction is with the C-terminal helical bundle (Fig. S1B). Rab2 is localised to the Golgi and was initially reported to act in traffic between the ER and Golgi (Cheung et al., 2002; Tisdale and Balch, 1996). However, in *C. elegans*, which lack FAM114A, Rab2 appears to be primarily involved in the formation of dense-core vesicles (Ailion et al., 2014; Sumakovic et al., 2009), and in *Drosophila* there is genetic evidence for roles in constitutive secretion, lysosome function and dense core vesicle production (Fujita et al., 2017; Götz et al., 2021; Ke et al., 2018; Lorincz et al., 2017). However, Rab2 binds to quite a wide range of effectors and so it may simply serve to recruit a diverse set of proteins to the correct part of the Golgi, and this set of proteins can vary between species. Finally, it should be noted that the tryptophan-glycine repeat motif conserved in all FAM114A proteins has properties similar to those of neutral amphipathic helixes that have been proposed to direct binding or proteins to lipid bilayers, and so it could potentially augment the action of Rab2 in targeting of the FAM114A proteins to membranes (Drin et al., 2007; Van Hilten et al., 2024).

There is clearly much that remains to be learnt about the in vivo role of FAM114A proteins, but our work clearly indicates that they have a role in Golgi function and appear to be new additions to the growing list of proteins that serve to allow COPI-coated vesicles to transport different cargo in different parts of the Golgi stack.

## MATERIALS AND METHODS

### Plasmids

For a list of the plasmids used in this study see Table S2. GFP-tagged cytoplasmic tail chimeras were generated as described previously (Welch et al., 2021). Plasmids designed to delete *FAM114A1* and *FAM114A2* were generated by annealing complementary oligonucleotide pairs encoding single gRNAs with BbsI-compatible overhangs and cloning them into the BbsI-digested bicistronic CRISPR-Cas9 mammalian expression vector pX458 (pSpCas9[BB]-2A-GFP). The coding sequence of *FAM114A1* and *FAM114A2* was fused to a c-terminal GAGA linker and a GST tag and cloned into the baculoviral expression vector pAcebac1 (Geneva Biotech) and the bacterial expression vector pOPC. *Drosophila* CG9590 was fused to a C-terminal GSGSGS linker and a GST tag and cloned into pOPC.

### Antibodies

For a list of the antibodies used in this study see Table S2. To raise an antibody against CG9590, GST-tagged CG9590 was produced in bacteria and affinity purified with glutathione beads (see below). Purified GST-tagged CG9590 was freeze dried before being used for 5 rounds of immunisations of rabbits over 2 months (Davids Biotechnologie, Regensburg Germany). Rabbit serum was depleted of GST-specific antibodies and subsequently affinity purified using the GST-tagged CG9590 antigen immobilised on beads, being eluted from beads using a low pH buffer and immediately neutralised on elution.

### Mammalian cell culture

U2OS (ATCC), HEK293T cells (ATCC) and FibroGRO Xeno-Free human foreskin fibroblasts (Merck, gift from Martin Lowe) were maintained in Dulbecco’s modified Eagle’s medium (DMEM, Thermo Fisher Scientific) with 10% fetal bovine serum (FBS, Thermo Fisher Scientific) and penicillin-streptomycin (PS) in a humidified incubator at 37°C with 5% CO_2_. Flp-In T-Rex HEK293 cell lines (Thermo Fisher Scientific) expressing mitochondrial-relocated golgin-BirA* fusions (from John Shin) were maintained in DMEM with 10% FBS, PS, 5 μg/mL blasticidin (Generon) and 100 μg/mL hygromycin (Thermo Fisher Scientific). Cells were passaged 1:10 every 3-4 days by trypsinisation and were regularly screened for mycoplasma contamination (Mycoalert, Lonza).

### Insect cell culture

D.Mel-2 cells were maintained in Schneider’s *Drosophila* medium (Thermo Fisher Scientific) with 10% FBS and antibiotics at 24°C. Cells were subcultured at a ratio of 1:10 every 3-4 days by detaching cells through tapping of the flask and the cell suspension diluted in fresh medium in a fresh flask. Large scale D.Mel-2 cultures were prepared by diluting cells to a density of 10^6^ cells/mL in Insect Xpress culture medium (Lonza) in Erlenmeyer flasks. Cells were incubated at 25°C with shaking at 140 rpm and subcultured by dilution at a ratio of 1:10 every 3-4 days.

GST-tagged FAM114A proteins were produced in Sf9 cells using the MultiBac baculoviral expression system (Geneva Biotech). Sf9 cells were seeded at a density of 10^6^ cells/cm^2^ in 6-well plates in Insect Xpress culture medium at 27°C and allowed to adhere for at least 10 minutes. Cells were transfected with 2 μg of bacmid DNA using Fugene HD transfection reagent (Promega) according to the manufacturer’s protocol. Cells were incubated for 3-5 days and the medium containing virus was used to inoculate a 50 mL culture of cells at a density of 2x10^6^ cells/ml in 250 mL Erlenmeyer flasks. Cells were cultured at 27°C with shaking at 140 rpm for 3-5 days. Cells were pelleted at 2500 x g for 10 minutes and the pellet stored on ice or snap frozen in liquid nitrogen prior to protein purification.

Alternatively, the supernatant containing the virus was used to inoculate larger cultures or to enhance the viral titre. The supernatant was preserved in the presence of 2% FBS at 4°C in darkness for medium term storage.

### Deletion of *FAM114A1* and *FAM114A2* by CRISPR-Cas9 gene editing

CRISPR-Cas9 gene editing was used to simultaneously knockout *FAM114A1* and *FAM114A2* in U2OS cells by targeting early constitutive exons with small out-of-frame deletions. Exon 3 of *FAM114A1* was simultaneously targeted at 5’-GTGCAGGGGCTGCCGCCATT-3’ and 5’ CCAACACCAGCTGACCCCAG-3’ and exon 2 of *FAM114A2* was targeted at 5’-ACTCTCTGGTTTGGCACCT-3’ and 5’-GGGGCTGCTTCAGTTAGCAG-3’. U2OS cells were seeded in T-75 flasks in culture medium and maintained in a humidified incubator with 5% CO2 at 37°C. Once cells reached 50-80% confluency they were transfected with CRISPR-Cas9 plasmids using polyethylenimine. 24 hours after transfection, single GFP-positive clones (i.e. cells expressing Cas9-2A-GFP) were sorted into 96-well plates (MoFlo Cell Sorter, Beckman Coulter). Candidate knockout clones were validated by immunoblot and the lead clone was further validated by mass spectrometry. *GOLPH3* and *GOLPH3L* were simultaneously deleted in the ΔΔ*FAM114A1*, *FAM114A2* U2OS background as described previously (Welch et al., 2021).

### PiggyBac transposon stable cell line generation

Stable cell lines expressing GFP-tagged cytoplasmic tail chimeras under a cumate-inducible promoter were generated by PiggyBac transposition (System Biosciences). Wild-type; ΔΔ*FAM114A1*, *FAM114A2* and ΔΔΔΔ*FAM114A1*, *FAM114A2*, *GOLPH3*, *GOLPH3L* CRISPR knockout U2OS cells were cultured to 50% confluency in 6-well plates and subsequently transfected with 0.2 μg PiggyBac transposase (PB210PA-1) and 0.5 μg of the PiggyBac-compatible expression plasmid. Cells were expanded to T-75 flasks 2 days after transfection and cells were subject to selection in culture medium with 0.5-1 μg/mL puromycin (Sigma) 3 days after transfection. Cells were cultured in selection media for several weeks until the polyclonal pool of integrants had reached confluency. Cell lines were immediately cryopreserved and maintained in selection medium containing 60 μg/mL cumate (System Biosciences) for at least one passage prior to assay.

### siRNA Knockdown of FAM114A in fibroblasts

Foreskin fibroblasts at 60-80% confluency 24 hours after seeding were treated with ON Targetplus SMARTpool siRNA oligos targeting *FAM114A1* and *FAM114A2* separately or simultaneously or were treated with a non-targeting negative control siRNA (Horizon Discovery) using Lipofectamine RNAiMAX transfection reagent according the manufacturer’s instructions (Thermo Fisher Scientific). Cells were treated with siRNA on day 1 and 3 after seeding, and on day 6 were washed once gently with PBS and lysed in plate with 1x LDS sample buffer with 10% TCEP. The lysate was sonicated and clarified by centrifugation before being subject to immunoblot analysis.

### MitoID proximity-dependent labelling assay

Doxycycline-inducible stable Flp-In T-REx HEK293 cell lines expressing mitochondrial-relocated golgin-BirA* fusion proteins were induced with 1 μg/ml doxycycline (Sigma) in culture medium once they reached approximately 80% confluency. 24 hours after induction, cells were treated with 0.5 μM nocodazole (Sigma), 50 μΜ biotin (Sigma) and 1 μg/mL doxycycline in culture medium for a further 9 hours. Cells were harvested and lysed for a streptavidin pulldown. Dynabeads One Streptavidin T1 beads (Thermo Fisher Scientific) were washed once in lysis buffer (50 mM Tris HCl pH 7.4, 150 mM NaCl, 1 mM EDTA, 0.5% Triton X-100, 1 mM PMSF, 1x cOmplete EDTA-free protease inhibitor (COMP)) using a DynaMag-2 magnetic stand (Thermo Fisher Scientific). Cell lysates were added to the washed beads and incubated overnight with agitation at 4°C. The beads were washed twice in wash buffer 1 (2x SDS with 1x COMP) for 8 minutes, 3 times in wash buffer 2 (50 mM Tris HCl pH 7.4, 500 mM NaCl, 1 mM EDTA, 1% Triton X-100, 0.1% deoxycholate, 1x COMP) for 8 minutes and 3 times in wash buffer 3 (50 mM Tris HCl pH 7.4, 50 mM NaCl, 1x COMP) for 8 minutes. Proteins were eluted by boiling at 98°C in 1x LDS, 10% β-mercaptoethanol and 6 mM biotin for 5 minutes. Samples were resolved by SDS PAGE, and gel slices were sent for mass spectrometry analysis.

### Cell lysis

Pelleted mammalian, bacterial and insect cells were resuspended in lysis buffer (as described for the streptavidin pulldown). Large scale cultures were sonicated on ice for 1 minute with 10 second on-off cycles (bacteria cells) or for only 10 seconds (mammalian and insect cells) using a lance sonicator (Sonic Vibra-Cell, 45% amplitude). Smaller scale cultures were sonicated using a water sonicator for 1 minute (mammalian and insect cells) or 3 minutes (bacteria cells) with 10 second on-off cycles (Misonix 300, amplitude 5.0).

Lysates were immediately placed on fresh ice for at least 5 minutes to mitigate heat generation from sonication. Lysates were subject to agitation at 4°C for a further 10 minutes prior to clarification by centrifugation at 16,000-32,000 x g for 10 minutes at 4°C. Where required, protein content was quantified using the Pierce BCA Protein Assay Kit and lysates normalised. Protein samples were kept on ice prior to downstream purification or were resolved by SDS PAGE.

### GST affinity chromatography

Glutathione Sepharose 4B beads (GE Life Sciences) were equilibrated in lysis buffer prior to pelleting at 100 x g for 1 minute and removal of the supernatant. Lysates containing GST-tagged fusion proteins were incubated with beads for 30 minutes with agitation at 4°C. Beads were then washed once with lysis buffer with 150 mM NaCl, once with lysis buffer with 500 mM NaCl and then another four times with lysis buffer with 150 mM NaCl. For the purification of GST-tagged CG9590 for rabbit immunisations, the fusion protein was eluted in buffer consisting of 50 mM Tris HCl with 25 mM reduced glutathione. For pulldowns upstream of mass spectrometry analysis, lysates containing prey proteins were preincubated on glutathione Sepharose beads at 4°C for 30 minutes to preclear non-specific interactors. Prey lysates were mixed with beads loaded with GST-fusion baits and the mixtures were incubated at 4°C for 1 hour with agitation. Beads washed 5 times in lysis buffer prior to elution in lysis buffer with 1.5 M NaCl. The prey proteins in the eluate precipitated with TCA/acetone and resolubilised in 1x LDS with 10% β-mercaptoethanol or TCEP. Bait proteins were eluted by boiling in 2x LDS with 10% β-mercaptoethanol or TCEP.

### Lectin labelling of cells

Wild-type and knockout U2OS cell lines were seeded at 2x10^4^ cells/cm^2^ in T-75 flasks in culture medium at 37°C with 5% CO2. At 80-90% confluency, cells were washed in EDTA solution and detached using Accutase (Sigma) for 2 minutes at 37°C. Cells were resuspended in ice cold FACS buffer (2% FBS in PBS) and approximately 1 million cells were transferred to round-bottom 96-well plates. Cells were pelleted by centrifugation at 300 x g for 5 minutes, the supernatant removed, and cells washed by resuspension in FACS buffer. Cells were stained with fluorescein-labelled lectins at 20 μg/mL (Vector Biolabs) and a fixable eFluor 780 viability dye diluted 1:1000 (Thermo Fisher Scientific) in FACS buffer on ice in darkness for 30 minutes. Non-specific binding was controlled by preincubation of the lectin with saturating concentrations of competing sugars at least 30 minutes prior to addition to cells. Finally, cells were washed 3 times in FACS buffer, fixed in 4% paraformaldehyde (PFA) diluted in PBS for 20 minutes at room temperature and washed a further two times in FACS buffer. Suspensions were kept at 4°C in darkness until required and were filtered using a 100 μm plate filter prior to loading on an LSRII flow cytometer (BD Biosciences). Gates were applied and density curves generated using FlowJo V10. Briefly, singlets were gated based on forward and side scatter, dead cells were excluded from analysis using the viability dye.

### Flow cytometry Golgi retention assay

Inducible stable cell lines expressing GFP-tagged cytoplasmic tail chimeric reporters were cultured in 6-well plate format in selection media containing 60 μg/mL cumate for at least a week prior to analysis. Once cells reached 80-90% confluency, they were washed once with EDTA solution and were detached from the plate in Accutase for 2 minutes at 37°C. Cells were resuspended in selection media and cell suspensions were transferred into a deep 96-well plate. Cells were pelleted at 300 x g for 5 minutes and were resuspended in ice cold FACS buffer. Suspensions were transferred to a round-bottomed 96-well plate and were resuspended in a cocktail consisting of an Alexa Fluor (AF) 647-conjugated anti-GFP antibody (BioLegend) and a eFluor 780 fixable viability dye diluted in FACS buffer. Cells were incubated on ice in darkness for 30 minutes before being washed, fixed and analysed as described for lectin stains.

### Mass Spectrometry

Protein samples generated from the MitoID assay and GST affinity chromatography were resolved by SDS PAGE and gels stained with InstantBlue Coomassie stain (Expedeon). Gel slices were excised for trypsin digestion and analysis by Nanoflow reverse-phase liquid chromatography-mass spectrometry using the Velos Orbitrap mass spectrometer (Thermo Fischer Scientific) as described previously (Gillingham et al., 2019). For spectral count analysis of the results of the MitoID assay, Mascot (Matrix Science) was used to search for peptides against the UniProt human proteome and further filtered using Scaffold (Proteome Software Inc). MitoID spectral counts were compared using D-score analysis from the open-source ComPASS platform (Sowa et al., 2009).

For whole cell proteomic analysis cells were lysed in 8 M urea with 20 mM Tris HCl before being sonicated and cleared by centrifugation. Total protein concentration was measured using a BCA assay (Pierce) and adjusted to 200 μg/mL. Protein samples were reduced with 5 mM DTT, alkylated with 10 mM iodoacetamide and subject to sequential protein digestion with Lys-C and trypsin (Promega). Digestion was halted with formic acid, precipitates cleared by centrifugation and supernatants desalted. Peptides were labelled using TMT 10plex reagent and separated on an offline HPLC. Finally, peptides were resolved on a 3000 RSLC Nano System (Thermo Fisher Scientific) and peptides were analysed via a nanospray ion source into a Q Exactive Plus hybrid quadrupole Orbitrap mass spectrometer (Thermo Fisher Scientific).

Mass spectrometry data generated from GST affinity chromatography and whole cell proteomic analysis was analysed using MaxQuant and peptides were searched against the UniProt human or *Drosophila* proteome using Andromeda (Cox and Mann, 2008; Cox et al., 2011). The Perseus platform was used to filter samples and to convert protein LFQ intensities to logarithmic values (Tyanova et al., 2016). Missing values were imputed using the default settings, statistical tests made using Welch’s or student’s t-tests, and volcano plots generated.

### Immunofluorescence of tissue culture cells

U2OS cells were trypsinised, added to complete media, seeded onto microscope slides (Hendley-Essex), and incubated at 37°C with 5% CO2. The next day cells were washed in PBS, fixed in 4% PFA in PBS for 20 minutes at room temperature and washed again in PBS. Cells were permeabilised in 10% Triton X-100 in PBS for 10 minutes and detergent was removed with five PBS washes. Cells were blocked in 20% FBS with 1% Tween-20 in PBS for 1 hour, blocking buffer was removed and cells were incubated with antibody diluted in blocking buffer for an hour. Cells were washed first in PBS, then in blocking buffer and then they were incubated with an anti-rabbit AF555 secondary antibody (Thermo Fisher Scientific) and an AF488 GFP booster (Chromotek) diluted in blocking buffer for 1 hour. They were washed again in PBS, then in blocking buffer and finally in PBS before most liquid was aspirated, and cells were mounted in Vectashield (Vector Biolabs). Slides were imaged using a 63x oil-immersion objective on a Leica TCS SP8 confocal microscope.

### Fly Stocks

*Drosophila melanogaster* stocks and crosses were kept on Iberian food (5.5% (w/v) glucose, 3.5% (w/v) organic wheat flour, 5% (w/v) yeast, 0.75% (w/v) agar, 16.4mM Nipagin (methyl-4-ydroxybenzoate) and 0.004% (v/v) propionic acid) at 25°C and 50% relative humidity with a repeating 12h light/12h dark cycle. The following stocks were used: Oregon R as a control, CFD2_nos-Cas9 (Port et al., 2014), Df(3R)BSC569 (BDSC #25670) - a genomic deficiency that includes the *CG9590* locus, *fkh*-Gal4 on the second chromosome (BDSC # 78061), *FAM114A^Δ43^* (*CG9590* null mutant, this study), UAS-FAM114A-GFP (this study).

### Generation of *CG9590/FAM114A* null mutants

CRISPR/Cas9 was used to generate a *Drosophila CG9590* null mutant. To remove the entire coding region of *CG9590* a pair of gRNAs was chosen targeting either end of the genomic locus. Both were cloned separately into pCFD3 as previously described ((Port et al., 2014), http://www.crisprflydesign.org/)). pCFD3-gRNA-CG9590_2 and pCFD3-gRNA-CG9590_4 were then co-injected into CFD2_nos-Cas9 embryos at a concentration of 100 ng/µl each. G0 flies were crossed to balancer stocks and F1 males were used to set up single crosses to generate stable lines. Once crosses were going, males were removed from vials and used for diagnostic PCRs and sequencing. The genomic DNA was isolated using microLYSIS Plus (Clent Life Sciences). We recovered *FAM114A^Δ43^* that removes the entire *CG9590* genomic locus.

### Generation of UAS-FAM114-GFP stock

The *CG9590* cDNA and a C-terminal eGFP with a GHGTGSTGSGSSR linker in between were cloned into pUAS-K10attB using NEBuilder HiFi DNA Assembly (NEB). Briefly, UAS-K10attB was cut with NotI and XbaI and the *CG9590* cDNA and eGFP amplified by PCR with homology arms and the linker sequence added to the oligos for HiFi DNA assembly. The UAS-FAM114A-GFP construct was then injected into embryos carrying an attP40 landing site and expressing the phiC31 Integrase under the vasa promoter. Injections were performed by John Overton (Gurdon Institute, Cambridge, UK). Successful transformants were identified by the presence of red eyes and used to make stable lines.

### Immunofluorescence of *Drosophila* 3^rd^ instar larval salivary glands

Wandering 3^rd^ instar larvae were collected and salivary glands were dissected in PBS. Tissues were fixed in fresh 4% PFA for 30min and then permeabilised for 4x30min in PBS-0.3% Triton-X100 (Sigma). Tissues were then blocked for 4x30min in PBS-0.1% Triton X-100, 5% BSA (Cell Signaling Technology) and primary antibodies were incubated in PBS-0.1% Triton X-100, 5% BSA o/n at 4C. Tissues were washed 4x30min in PBS-0.1% Triton-X100 and secondary antibodies were incubated in PBS-0.1% Triton X-100, 5% BSA o/n at 4C. The Chromotek GFP-Booster Atto 488 (Proteintech) was used to boost the GFP signal. Tissues were washed 4x30min in PBT-0.1% Triton-X100 and equilibrated in Vectashield with DAPI (2BScientific) o/n at -20C. Tissues were then mounted in Vectashield with DAPI. Images were taken with a confocal microscope (Zeiss 710 or 900) and processed in Fiji. Primary and secondary antibodies are in Table S2.

### Immunoblotting of *Drosophila* samples

For each genotype five salivary gland pairs from wandering 3^rd^ instar larvae were dissected in PBS and immediately transferred into RIPA buffer (Sigma) plus protease inhibitors (cOmplete, Roche, PMSF, Sigma) on ice. Samples were homogenised using a Kimble pellet pestle (DWK Life Science) and left on ice for 25 min. NuPAGE 4xLDS sample buffer (Invitrogen) and 5% β-mercaptoethanol (Sigma) were added and the samples heated at 90°C for 10min. Samples were run on a NuPAGE 4-12% Bis-Tris mini gel (Invitrogen) using MES buffer (Formedium). After transfer to nitrocellulose (Amersham) the membrane was blocked in PBS, 0.1% Tween 20, 3% skimmed milk powder and 1% BSA for 1h at RT. The membrane was cut and primary antibodies were added o/n at 4°C in PBS, 0.1% Tween 20, 1% BSA, 3% skimmed milk powder (Marvel). Membranes were washed and secondary antibodies were added for one hour at RT. After washing the blot was developed using SuperSignal West Femto Maximum Sensitivity Substrate (Thermo Fisher). A BioRad Chemidoc MP imaging system (BioRad) was used to acquire images. Primary and secondary antibodies in Table S2.

### VVA and PNA staining of 3^rd^ instar larval salivary glands

Lectin staining was done as described previously (Tian et al., 2013). Briefly, tissues were fixed for 30 min in fresh 4% PFA at RT. VVA and PNA were conjugated with fluorescein (Vector Labs) and applied at 1 μg/ml, with or without competing sugar: N-acetyl-*D*-galactosamine (GalNAc, 0.3M, Sigma) for VVA and D(+)-galactose (0.3M, Formedium) for PNA. 6-9 salivary glands from different larvae were analysed for each genotype in 2-3 technical repeats. Images were taken on a Zeiss 710 confocal microscope and processed in Fiji.

## Acknowledgements

We thank Catherine Rabouille, Gunter Merdes, Jennifer Richens, John Kilmartin, John Shin, Kelly Ten Hagen, and Martin Lowe for reagents and advice. Mass spectrometry analysis was performed at the Biological Mass Spectrometry and Proteomics Facility of the MRC LMB. Funding was provided by the Medical Research Council, as part of United Kingdom Research and Innovation (also known as UK Research and Innovation) file reference number MC_U105178783.

---

**Supplementary Figure 1.**
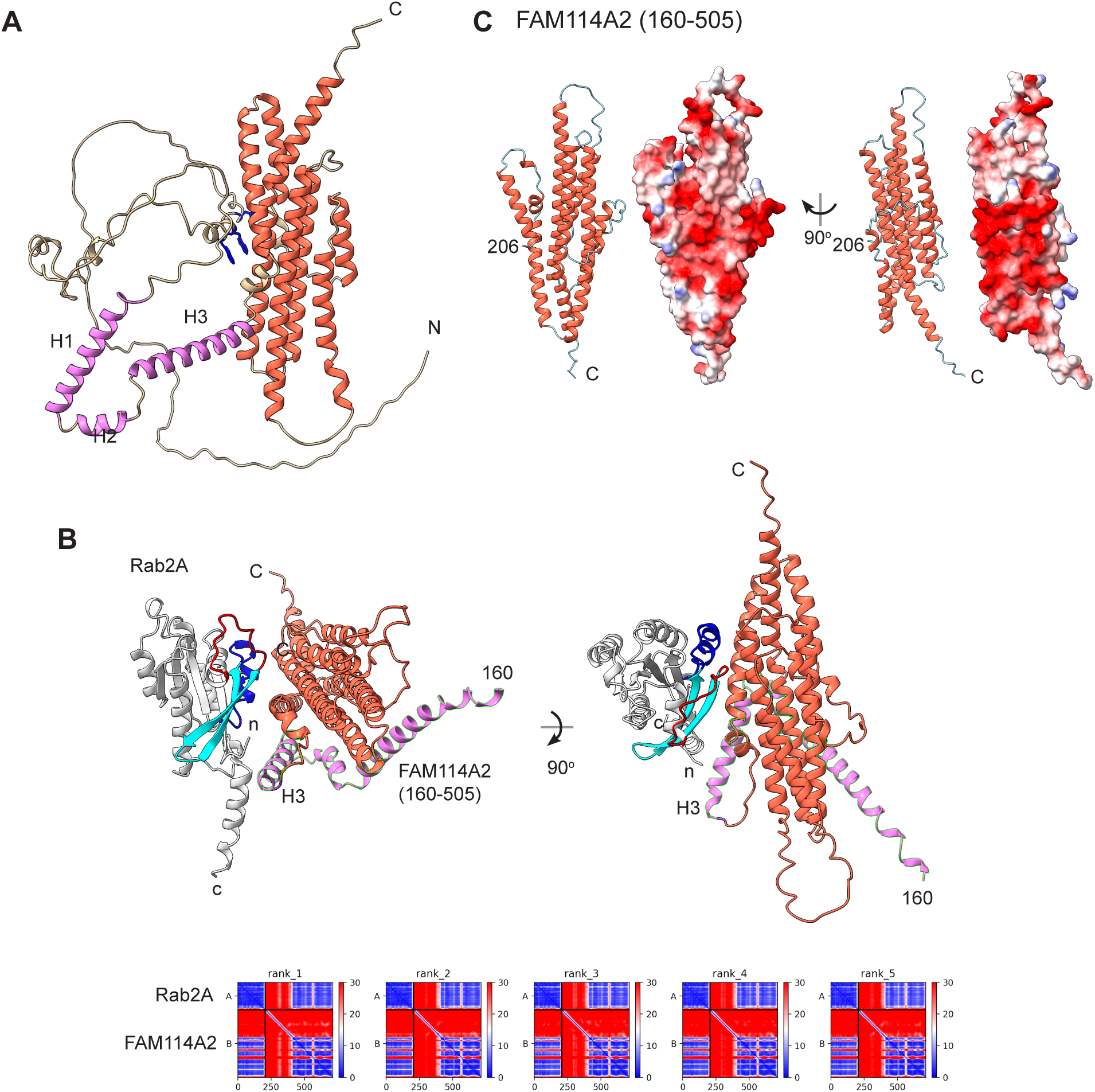
Predicted structures for FAM114A2. **(A)** AlphaFold prediction for the structure of human FAM11A2. From three recycles using mmseqs2_uniref as MSA mode and default settings in ColabFold. The C-terminal helical bundle in brick-red and preceding three helices in violet. The three tryptophans in the WG motif region are shown in blue. **(B)** AlphaFold prediction for the structure of human FAM11A2 in complex with Rab2A. From three recycles using mmseqs2_uniref as MSA mode and default settings in ColabFold, along with the PAE plot of all five models (lower scores are higher confidence and in blue). The interaction is predicted with high confidence (ipTM=0.893). For FAM114A2 the unstructured region (1-159) is omitted for clarity and colouring is as in (A). Rab2A is in grey with Switch-1 (red), interswitch (red) and Switch-2 (blue) indicated. As expected for a Rab:effector interaction, the Switch and interswitch regions make contact with FAM114A2 helical bundle. Helix 3 is predicted to also pack against Rab2A, but this has not been verified experimentally. **(C)** Electrostatic potential plot of the surface of the C-terminal helical bundle – red is negative, blue is positive. Two highly negative surfaces are present.

**Supplementary Figure 2.**
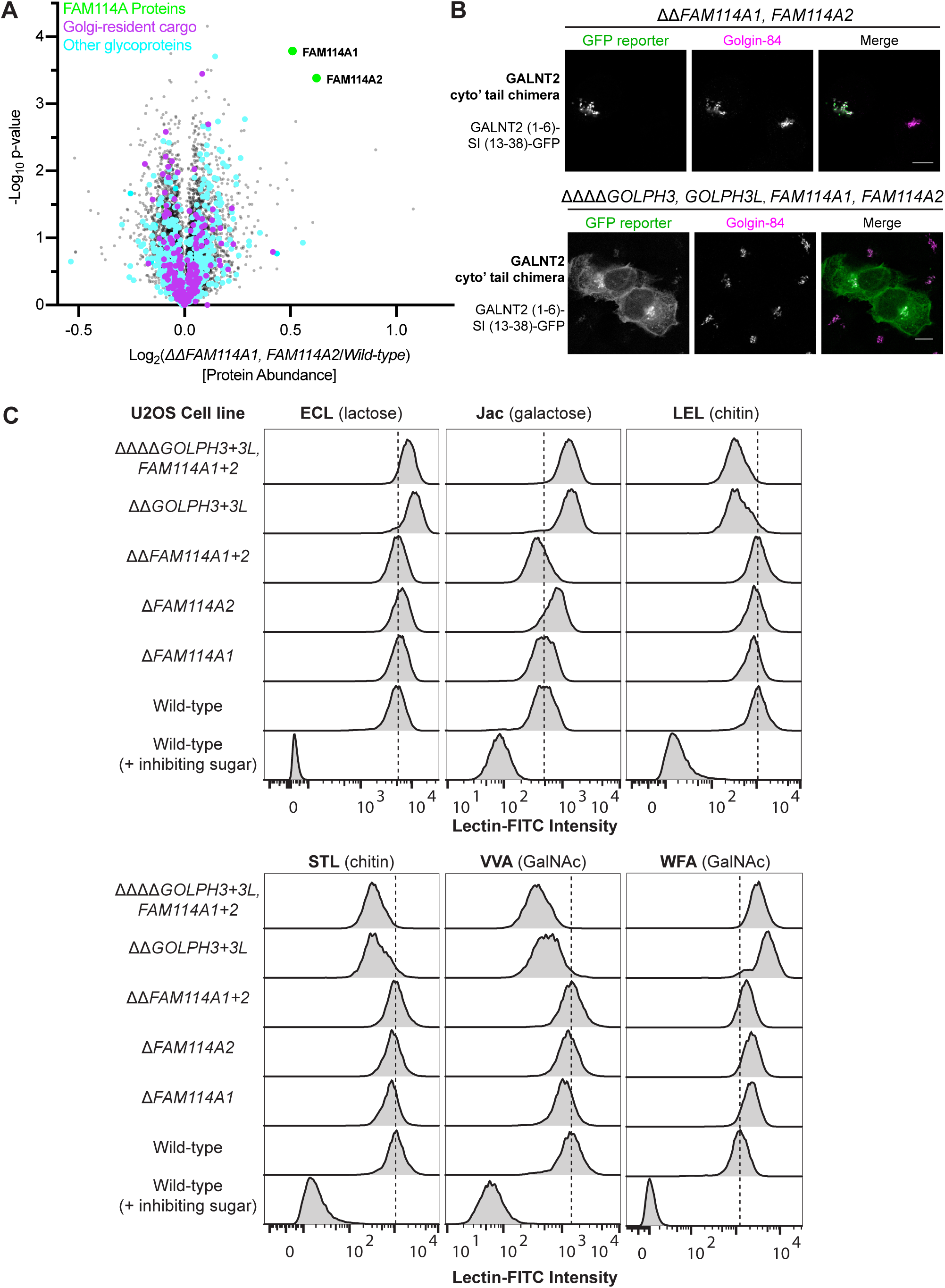
Analysis of cells lacking FAM114A1 and FAM114A2. **(A)** Volcano plot comparing the relative abundance of individual proteins in *ΔΔFAM114A1, FAM114A2* U2OS cells vs the wild-type parental control cell line. The Z-score was normalized based on the median and a Student’s t-test was applied to generate p-values. N=2. **(B)** Confocal micrographs showing a GFP-tagged cytoplasmic tail chimera expressed in *ΔΔFAM114A1, FAM114A2* and *ΔΔΔΔFAM114A1, FAM114A2, GOLPH3, GOLPH3L* U2OS cells. The GFP signal was enhanced with a GFP booster and golgin-84 was stained as a Golgi marker. **(C)** Density curves generated from flow cytometry analysis of different CRISPR knockout U2OS cell lines subjected to cell surface stains with a panel of different FITC-conjugated lectins (see top, in bold). Specificity of the lectin was validated in which cells were stained in the presence of saturating concentrations of a competing sugar (see top, brackets). Density curves are normalised to the mode value for each treatment. Dotted lines mark the mode intensity value for wild-type cells. At least 10,000 events were collected for each cell line. Singlets were gated for based on forward and side scatter, dead cells were excluded using a fixable viability stain. Shown are the results of a single repeat of a triplicate.

